# CHARMM-GUI Membrane Builder for Lipid Nanoparticles with Ionizable Cationic Lipids and PEGylated Lipids

**DOI:** 10.1101/2021.06.23.449544

**Authors:** Soohyung Park, Yeol Kyo Choi, Seonghoon Kim, Jumin Lee, Wonpil Im

## Abstract

A lipid nanoparticle (LNP) formulation is a state-of-the-art delivery system for genetic drugs such as DNA, mRNA, and siRNA, which is successfully applied to COVID-19 vaccines and gains tremendous interest in therapeutic applications. Despite its importance, a molecular-level understanding of the LNP structures and dynamics is still lacking, which makes a rational LNP design almost impossible. In this work, we present an extension of CHARMM-GUI *Membrane Builder* to model and simulate all-atom LNPs with various (ionizable) cationic lipids and PEGylated lipids (PEG-lipids). These new lipid types can be mixed with any existing lipid types with or without a biomolecule of interest, and the generated systems can be simulated using various molecular dynamics engines. As a first illustration, we considered model LNP membranes with DLin-KC2-DMA (KC2) or DLin-MC3-DMA (MC3) without PEG-lipids. The results from these model membranes are consistent with those from the two previous studies albeit with mild accumulation of neutral MC3 in the bilayer center. To demonstrate *Membrane Builder*’s capability of building a realistic LNP patch, we generated KC2- or MC3-containing LNP membranes with high concentrations of cholesterol and ionizable cationic lipids together with 2 mol% PEG-lipids. We observe that PEG-chains are flexible, which can be more preferentially extended laterally in the presence of cationic lipids due to the attractive interactions between their head groups and PEG oxygen. The presence of PEG-lipids also relaxes the lateral packing in LNP membranes, and the area compressibility modulus (*K*_A_) of LNP membranes with cationic lipids fit into typical *K*_A_ of fluid-phase membranes. Interestingly, the interactions between PEG oxygen and head group of ionizable cationic lipids induce a negative curvature. We hope that this LNP capability in *Membrane Builder* can be useful to better characterize various LNPs with or without genetic drugs for a rational LNP design.

## Introduction

A lipid nanoparticle (LNP) formulation represents a viable delivery system of genetic drugs such as DNA, mRNA, and siRNA for expressing a beneficial protein, silencing a pathological gene, or editing defective genes to treat various diseases, including cancer, hereditary disorders, and viral infections.^1,2^ A key advance in LNP formulations for effective delivery of genetic drugs to the cytoplasm of target cells has been identification and incorporation of optimized ionizable cationic lipids (**Fig. 1**) with apparent pKa values around 6.5,^3^ such as dilinoleyl-methyl-4-dimethylaminobutyrate (DLin-MC3-DMA or MC3)^3^ or dilinoleyl-4-dimethylaminoethyl-[1,3]-dioxolane (DLin-KC2-DMA or KC2).^4^ In popular ethanol-loading LNP formulations, these cationic lipids whose main role is entrapment of nucleic acid polymers at low pH (∼4) are mixed with so-called helper lipids such as cholesterol (CHOL), phospholipids, and PEGylated lipids (in short, PEG-lipids).^5^ Clearly, LNP-mRNA formulations have become mankind’s hope for COVID-19 vaccines.^6^

**Fig. 1.**
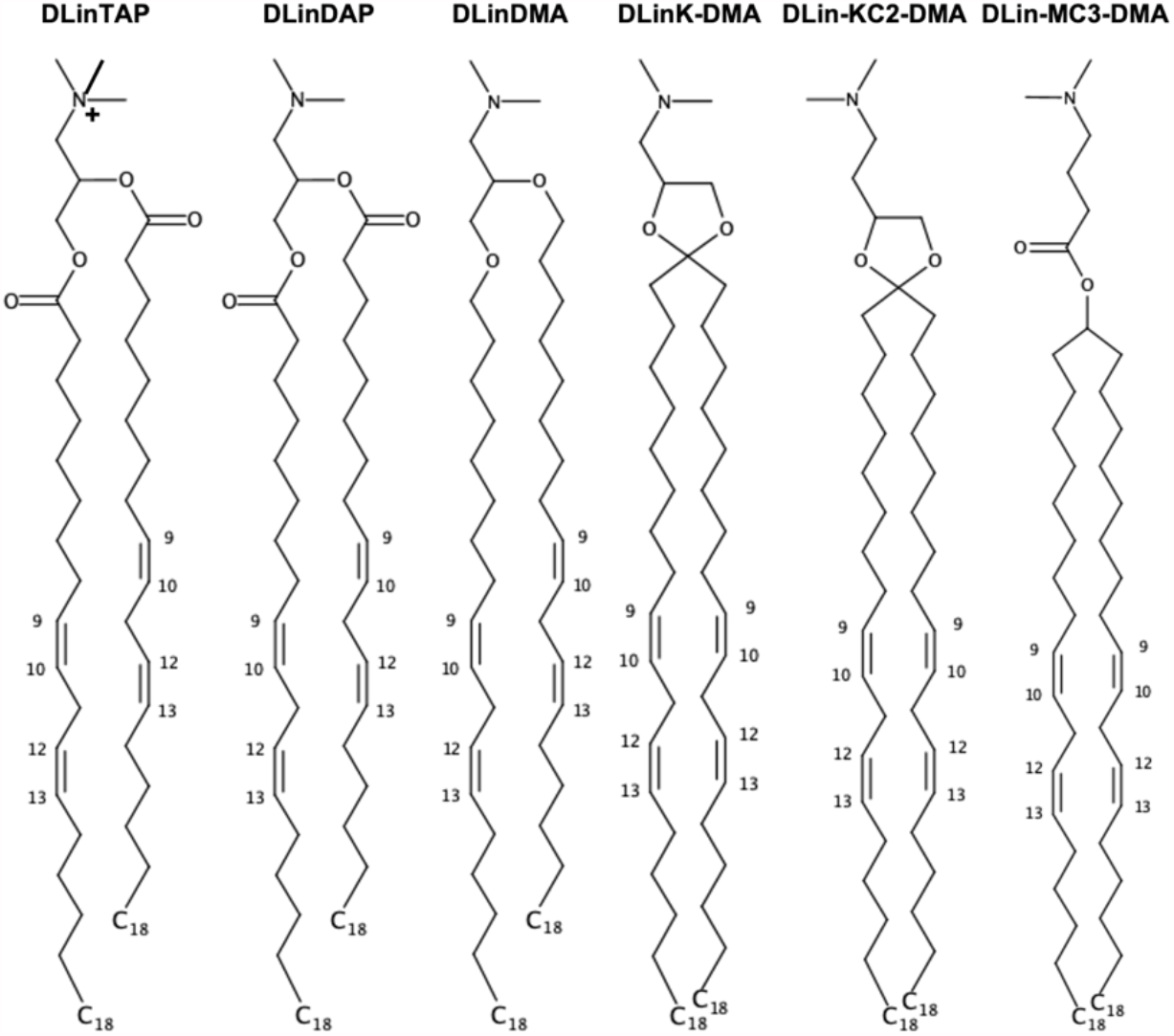
Chemical structures of selected (ionizable) cationic lipids available in Membrane Builder. 1,2-dilineoyl-3-trimethylammonium propane (DLinTAP), 1,2-dilinoleoyl-3-dimethylammonium propane (DLinDAP), 1,2-dilinoleyloxy-3-N,N-dimethylaminopropane (DLinDMA), 1,2-dilinoleyloxy-keto-N,N-dimethyl-3-aminopropane (DLinK-DMA or KC1), 1,2-dilinoleyl-4-dimethylaminoethyl-[1,3]-dioxolane (DLin-KC2-DMA or KC2), and dilinoleylmethyl-4-dimethylaminobutyrate (DLin-MC3-DMA or MC3).

A molecular-level understanding of the LNP structures and dynamics resulting from delicate interactions among constituent lipids and genetic drugs could provide insight into a rational LNP design. Surprisingly, to the best of our knowledge, there are not many all-atom modeling and simulation studies of these complex LNPs or their patches, probably due to difficulties in building such LNP systems with constituent lipids including various ionizable cationic lipids; n.b., all-atom models and simulations are also necessary to develop coarse-grained models for larger systems and longer simulations.^7,8^ Ramezanpour et al performed all-atom molecular dynamics (MD) simulations of neutral and cationic KC2 in palmitoyl-oleoyl-phosphatidylcholine (POPC) bilayers with or without CHOL.^9^ The main observation was that neutral KC2s at 10, 20, 30 mol% segregated at the bilayer center, whereas cationic KC2 head groups stayed at the bilayer-water interface. More recently, Ermilova and Swenson performed all-atom MD simulations of neutral MC3 in dioleoyl-phosphatidylcholine (DOPC) or dioleoyl-phosphatidylethanolamine (DOPE) bilayers.^10^ The main observation was that neutral MC3 head groups at 5 and 15 mol% positioned at the DOPE bilayer-water interface due to strong interactions between DOPE head groups and MC3 head groups and tails, whereas there are no such interactions between MC3 and DOPC. Note that these systems are relatively simple in that LNP formulations generally contain very high concentrations of CHOL and ionizable cation lipids together with 1-5 mol% PEG-lipids.^1,2^This work represents an important extension of CHARMM-GUI *Membrane Builder*^11,12^ to model and simulate all-atom LNPs with various (ionizable) cationic lipids and PEG-lipids. We have added 55 (ionizable) cationic lipids with 6 head groups of neutral and cationic forms and 5 tails, as well as PEG-lipids with any number of PEG units and with 20 PE-based and 20 diacylglycerol (DAG)-based tails. These new lipid types can be mixed with any existing lipid types with or without a biomolecule of interest, and the generated MD systems can be simulated using various programs, such as CHARMM,^13^ GROMACS,^14^ NAMD,^15^ LAMMPS,^16^ AMBER,^17^ GENESIS,^18^ OpenMM,^19^and Desmond.^20^ As a first illustration, we have built and simulated all systems of the aforementioned two simulation studies.^9,10^ In addition, to demonstrate *Membrane Builder’s* capability to build a realistic LNP patch, we built and simulated KC2-or MC3-containing LNP membranes with high concentration of CHOL (47 mol%) and ionizable cationic lipids (40 mol%) together with 2 mol% PEG-lipids.

## METHODS

### *Membrane Builder* implementation

For LNPs, *Membrane Builder* supports 6 cationic head groups (TAP, DAP, DMA, KC1, KC2, and MC3 in **Fig. 1**) with 5 lipid tails (dimyristoyl (14:0 / 14:0), dipalmitoyl (16:0 / 16:0), dioleoyl (18:1 / 18:1), distearoyl (18:0 / 18:0), and dilinoleoyl (18:2 / 18:2) in **Fig. 2a**). Except TAP, both neutral and cationic head group forms are available; e.g., DLKC2 is a neutral form and DLCK2H is a cationic form. Thus, the combination of head groups and tails yields a total of 55 (ionizable) cationic lipid types. A surface area of 60 Å^2^ is assigned to each (ionizable) cationic lipid as a tentative value and one can change it for their specific system building purpose. As shown in **Fig. 2a**, one can control the number of lipids in each leaflet using ratios or exact numbers.

**Fig. 2.**
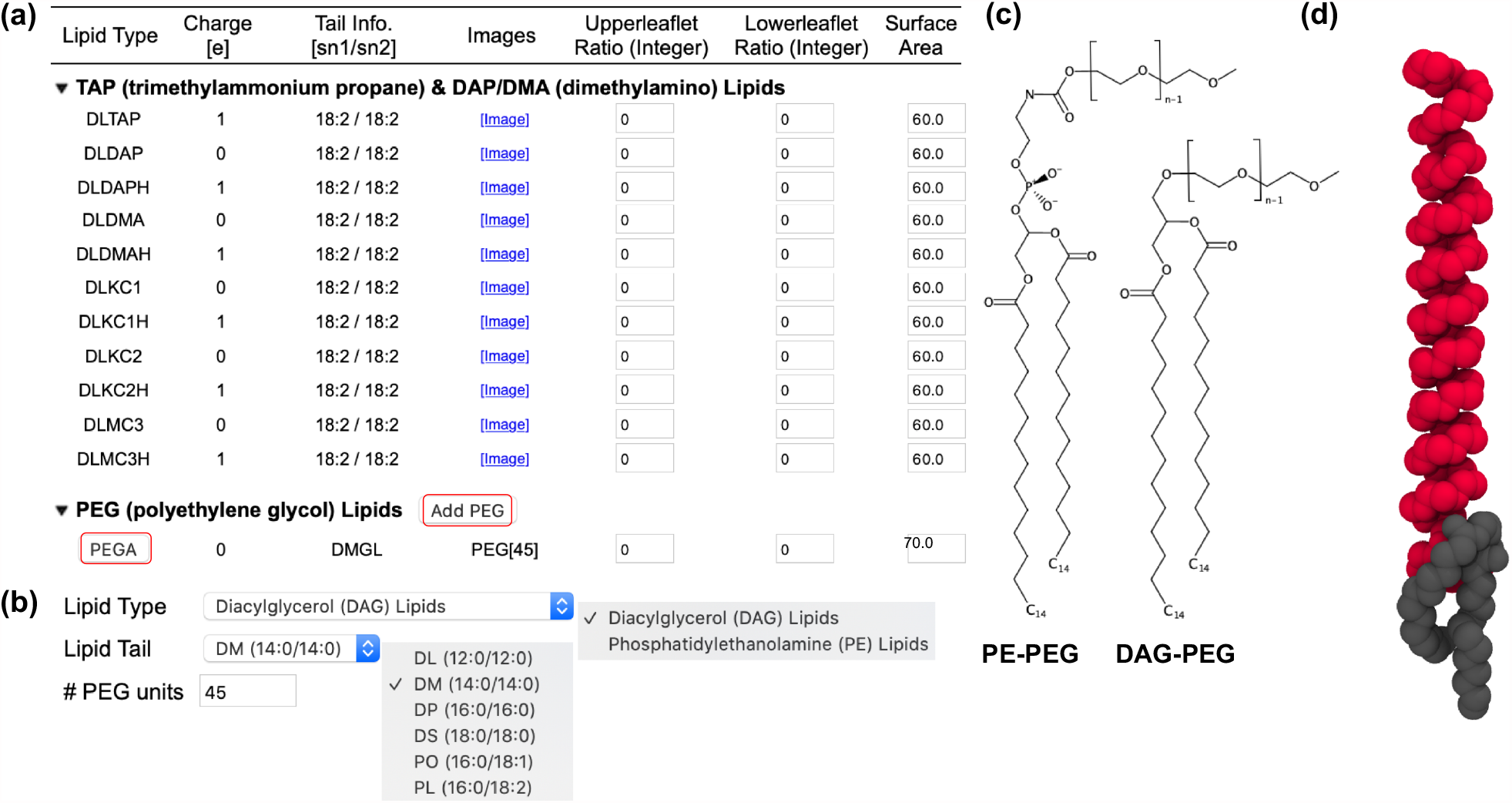
Illustrative snapshots of lipid selection in Membrane Builder for LNPs. (a) Two categories showing (ionizable) cationic lipids and PEG-lipids. Only cationic lipids with dilinoleoyl (18:2 / 18:2) tails are shown, although dimyristoyl (14:0 / 14:0), dipalmitoyl (16:0 / 16:0), dioleoyl (18:1 / 18:1), and distearoyl (18:0 / 18:0) tails are available for all head group types. Except TAP, both neutral and cationic head group forms are available; e.g., DLKC2 is a neutral form and DLCK2H is a cationic form. For PEG-lipids, one can click “PEGA” to select a lipid type and a tail with the options shown in (b) where DMG-PEG 2000 is constructed based on diacylglycerol (DAG) with dimyristoyl tail and 45 PEG units. Note that only 6 acyl tails are shown in (b) among 20 available tails. (c) Chemical structures of PE-and DAG-based PEG-lipids with n numbers of PEG units (d) An initial conformation of DMG-PEG 2000 that is used for the replacement method. The lipid tail and PEG part are represented by gray and red, respectively. Hydrogen atoms are omitted for clarity.

For PEG-lipids, we followed the workflow used for glycolipid and lipopolysaccharide structure generation in CHARMM-GUI.^21^ In this workflow, to define a PEG-lipid, one can click “PEGA” (in **Fig. 2a**) to select a lipid type and a tail with the options shown in **Fig. 2b**. Currently, *Membrane Builder* supports PE-based and DAG-based PEG-lipids (**Fig. 2c**) with 20 different acyl tails. **Fig. 2b** shows how to build DMG-PEG 2000 that is denoted as PEG[45] in **Fig. 2a**. One can add various PEG-lipids with different lipid types, tails, and numbers of PEG units by clicking “Add PEG” in **Fig. 2a**. Similar to other lipid types, one can control the number of lipids in each leaflet using ratios or exact numbers.

Unlike (ionizable) cationic lipids for which we can pre-generate a conformational library of 2,000 structures for the replacement method,^22,23^ it is impossible to prepare such a conformational library for each PEG-lipid molecule because lipid types, tails, and numbers of PEG units can be varied depending on user selection. Therefore, an additional step was introduced to generate a structure for each user-specified PEG lipid, and the generated structure is then used to replace the lipid-like pseudo spheres in the replacement step. PEG-lipids have very long flexible PEG-chains, and these chains can cause severe steric crashes with the neighboring PEG or lipid molecules during the system generation. In addition, an extended structure of DMG-PEG 2000 (with 45 PEG units) is about 190 Å long along the membrane normal (i.e., the *Z* axis), which requires an unnecessarily large system size. To avoid these problems, each PEG structure is generated to have a π-helical conformation along the Z axis. Then, additional procedures are introduced to relax the lipid head and tail portions with the PEG structure fixed in a π-helical conformation by performing a simulation with planar and cylindrical restraints; the former is to place the head group and terminal tail atoms in appropriate positions along the Z axis, and the latter is to make the entire PEG-lipid cylindrical along the Z axis. **Fig. 2d** shows an initial structure of DMG-PEG 2000 whose length is about 60 Å. Note that the building process is fast and does not depend on the number of PEG units.

#### Simulation systems and details

As a first illustration of our building protocol in *Membrane Builder*, we built and simulated the systems studies by Ramezanpour et al^9^ (Case 1 in **Table 1**) and Ermilova and Swenson^10^ using POPC and POPE instead of DOPC and DOPE (Case 2) to use the same lipid tails as in Case 1. Then, to further illustrate the capability of *Membrane Builder*, we built and simulated more realistic LNPs using KC2 (Case 3) and MC3 (Case 4). Here, we consider low and high pHs apart from pKa of KC2 and MC3, so each system has either a neutral or cationic form. All membrane systems are symmetric (i.e., the same compositions in both leaflets) and their membrane area is about 140 Å x 140 Å in the *XY* plane. Each system was neutralized with 150 mM NaCl (with counterions for KC2H and MC3H) using TIP3P water model.^24,25^ All (ionizable) cationic lipids and PEG molecules are based on the CHARMM36 lipid force field^26^ and the CGenFF.^27^

**Table 1.** Simulation system information

| Case 1 | KC2 | KC2H | POPC | CHOL |  | XYZ | #Na / #Cl | #atoms |
| --- | --- | --- | --- | --- | --- | --- | --- | --- |
| KC2-PC90<sup>a</sup> | 30 |  | 270 |  |  | 142 x 142 x 85 | 70 / 70 | 159,980 |
| KC2-PC80<sup>b</sup> | 59 |  | 236 |  |  | 140 x 140 x 85 | 67 / 67 | 155,769 |
| KC2-PC70<sup>c</sup> | 90 |  | 210 |  |  | 140 x 140 x 85 | 67 / 67 | 156,536 |
| KC2-PC-CH<sup>d</sup> | 114 |  | 95 | 171 |  | 142 x 142 x 85 | 73 / 73 | 160,822 |
| KC2H-PC90<sup>a</sup> |  | 30 | 270 |  |  | 142 x 142 x 85 | 70 / 130 |  |
| KC2H-PC80<sup>b</sup> |  | 59 | 236 |  |  | 140 x 140 x 85 | 67 / 185 | 155,594 |
| KC2H-PC70<sup>c</sup> |  | 90 | 210 |  |  | 140 x 140 x 85 | 67 / 247 | 156,728 |
| KC2H-PC-CH<sup>d</sup> |  | 114 | 95 | 171 |  | 142 x 142 x 85 | 72 / 300 | 161,210 |
| Case 2 | MC3 | MC3H | POPC | POPE |  | XYZ | #Na / #Cl | #atoms |
| MC3-PC95<sup>e</sup> | 15 |  | 285 |  |  | 143 x 143 x 85 | 70 / 70 | 160,652 |
| MC3-PC85<sup>f</sup> | 45 |  | 255 |  |  | 142 x 142 x 85 | 69 / 69 | 158,871 |
| MC3-PE95<sup>e</sup> | 17 |  |  | 323 |  | 141 x 141 x 85 | 69 / 69 | 164,977 |
| MC3-PE85<sup>f</sup> | 51 |  |  | 289 |  | 142 x 142 x 85 | 69 / 69 | 164,845 |
| MC3H-PC95<sup>e</sup> |  | 15 | 285 |  |  | 143 x 143 x 85 | 70 / 100 | 160,604 |
| MC3H-PC85<sup>f</sup> |  | 45 | 255 |  |  | 142 x 142 x 85 | 69 / 159 | 159,090 |
| MC3H-PE95<sup>e</sup> |  | 17 |  | 323 |  | 141 x 141 x 85 | 69 / 103 | 164,961 |
| MC3H-PE85<sup>f</sup> |  | 51 |  | 289 |  | 142 x 142 x 85 | 69 / 171 | 164,953 |
| Case 3 | KC2 | KC2H | DSPC | CHOL | PEGL<sup>i</sup> | XYZ | #Na / #Cl | #atoms |
| KC2-LNP<sup>g</sup> | 160 |  | 44 | 188 | 8 | 143 x 143 x 160 | 205 / 205 | 278,886 |
| KC2H-LNP<sup>g</sup> |  | 160 | 44 | 188 | 8 | 143 x 143 x 156 | 196 / 516 | 297,466 |
| KC2-DS-CH<sup>h</sup> | 160 |  | 52 | 188 |  | 143 x 143 x 85 | 72 / 72 | 164,192 |
| KC2H-DS-CH<sup>h</sup> |  | 160 | 52 | 188 |  | 143 x 143 x 85 | 72 / 392 | 164,349 |
| Case 4 | MC3 | MC3H | DSPC | CHOL | PEGL<sup>i</sup> | XYZ | #Na / #Cl | #atoms |
| MC3-LNP<sup>g</sup> | 160 |  | 44 | 188 | 8 | 143 x 143 x 141 | 169 / 169 | 246,003 |
| MC3H-LNP<sup>g</sup> |  | 160 | 44 | 188 | 8 | 143 x 143 x 145 | 176 / 496 | 250,233 |
| MC3-DS-CH<sup>h</sup> | 160 |  | 52 | 188 |  | 143 x 143 x 85 | 72 / 72 | 163,730 |
| MC3H-DS-CH<sup>h</sup> |  | 160 | 52 | 188 |  | 143 x 143 x 85 | 72 / 392 | 164,580 |

<sup>a</sup>KC2 (or KC2H) : POPC = 1 : 9. <sup>b</sup>KC2 (or KC2H) : POPC = 2 : 8. <sup>c</sup>KC2 (or KC2H) : POPC = 3 : 7. <sup>d</sup>KC2 (or KC2H) : POPC : CHOL = 30 : 25 : 45. <sup>e</sup>MC3 (or MC3H) : POPC (or POPE) = 5 : 95. <sup>f</sup>MC3 (or MC3H) : POPC (or POPE) = 15 : 85. <sup>g</sup>KC2 (or KC2H or MC3 or MC3H) : DSPC : CHOL : PEGL = 40 : 11 : 47 : 2. <sup>h</sup>KC2 (or KC2H or MC3 or MC3H) : DSPC : CHOL = 40 : 13 : 47. <sup>i</sup>DMG-PEG 2000.

All simulations were performed using OpenMM^19^ with inputs generated by CHARMM-GUI.^28^ Following *Membrane Builder*’s default six-step equilibration protocol,^22,23^ NVT (constant particle number, volume, and temperature) dynamics was first applied with a 1-femtosecond (fs) time step for 250 picoseconds (ps). Subsequently, the NPT (constant particle number, pressure, and temperature) ensemble was applied with a 1-fs time step (for 125 ps) and with a 2-fs time step (for 1.5 nanoseconds (ns)). During the equilibration, positional and dihedral restraint potentials were applied to lipid and water molecules, and their force constants were gradually reduced. A production run was performed for 2 microseconds (μs) (for KC2-LNP, KC2-DS-CH, MC3-LNP, and MC3-DS-CH in **Table 1**) or 1.5 μs (for all other systems) with a 4-fs time step using the hydrogen mass repartitioning technique^29,30^ without any restraint potential; see **Fig. S1** in ESI† for the time series of the membrane area of each system. The SHAKE algorithm was applied to the bonds containing hydrogen atoms.^31^ The van der Waals interactions were cut off at 12 Å with a force-switching function between 10 and 12 Å,^32^ and the electrostatic interactions were calculated by the particle-mesh Ewald method.^33^ The temperature (at 310 K) and the pressure (at 1 bar) were controlled by Langevin dynamics with a friction coefficient of 1 ps^−1^ and a semi-isotropic Monte Carlo barostat, respectively.^34,35^ **Fig. 3** shows system images for KC2-PC-CH, MC3-PE85, and MC3-LNP.

**Fig. 3.**
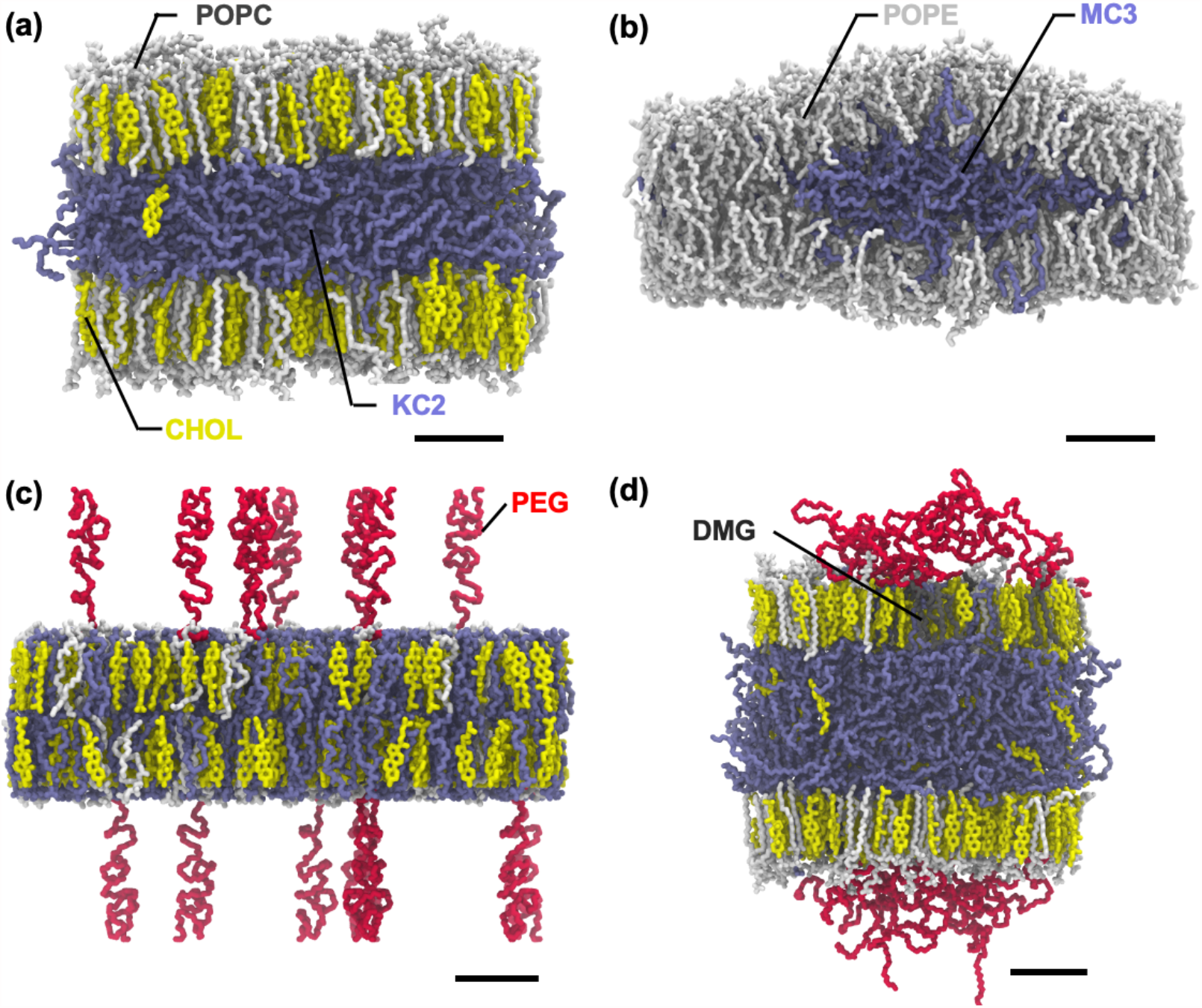
Snapshots of (a) final KC2-PC-CH system, (b) final MC3-PE85 system, (c) initially relaxed MC3-LNP system, and (d) final MC3-LNP system. Phospholipids (POPC, POPE, and DSPC), CHOL, and ionizable cationic lipids (KC2 and MC3) are shown in sticks. Each component is shown in different color: phospholipids in white, CHOL in yellow, KC2 and MC3 in ice-blue, PEG-lipid tail in grey, and PEG-chain in red. For clarity, water and ions are omitted and only heavy atoms are shown. The full name of each lipid type is given in the main text. The scale bar represents 20 Å.

#### Analysis

The last 600-ns trajectories from each simulation were used for analysis. From the time series of each system size, its *XY*-area (*A*) was calculated (**Fig. S1** in ESI†). The lateral packing of bilayers was analyzed by calculating the area compressibility modulus (*K*_*A*_) and deuterium order parameters (*S*_CD_). *K*_*A*_ is defined as *K*_*A*_ = *k*_B_*T* ⟨A⟩ / ⟨d*A*_2_⟩, where *k*_B_ is the Boltzmann constant, *T* is temperature, and <*A*> and <*dA*^2^> are the mean area and its mean square fluctuation, respectively.*S*_CD_ are defined as

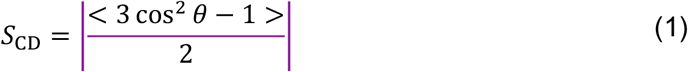

where *θ* is the angle between a C-H bond vector and the membrane normal, and the bracket represents the time and ensemble average. The bilayer thickness (*d*_B_) was defined as the distance between the average *Z*-positions of P atoms in two leaflets. The distributions of components (phospholipid, ionizable lipid, CHOL, PEG-lipids, ions, and water) along the membrane normal were analyzed by calculating their number density distribution, where the bilayer was recentered to be located at *Z* = 0.

The torque density of monolayer (𝒯) is the first moment of the lateral pressure profile, *p*(*Z*) = *p*_.T_(*Z*) – *p*_N_(*Z*), where *p*_T_(*Z*) = [*p*_*xx*_(*Z*)+*p*_*yy*_(*Z*)]/2 and *p*N(*Z*) = *pzz*(*Z*) = *p*N are the tangential and normal components of pressure tensors to the membrane surface (see **Lateral pressure profile and torque density** in ESI†). To calculate *p*(*Z*), the velocity (and associated coordinate) trajectories were generated from the restart files for the last 600 ns, which were saved every 0.1 ns. Two in-house python classes VELFile and VELReporter were used for velocity trajectories, which were modified from DCDFile and DCDReporter for generation of coordinate trajectories in OpenMM. From the (recentered coordinate and velocity) trajectories, *p*(*Z*) was calculated using an in-house version of CHARMM, where Harasima contour of slab geometry^36^ was employed for calculation of *p*_T_(*z*). Because *p*_N_(*Z*) cannot be correctly estimated by Harasima contour,^37^ with an assumption of a tensionless bilayer, ∫ *dZ* [*p*_$_(*Z*) - *p*_%_] = 0, *p*_N_ was calculated by *p*_%_ = ∫ *dZ p*_$_(*Z*)/*L*_&_, where *L*_z_ is the *Z*-dimension of the system.

The PEG conformation was analyzed by the end-to-end distance (⟨*h*^2^⟩^1/2^), its *X*-, *Y*-, *Z*-components (⟨*h*_*x*_^2^⟩^1/2^, ⟨*h*_*y*_^2^⟩^1/2^, ⟨*h*_*z*_^2^⟩^1/2^), and its thickness (*d*_PEG_) for which PEG oxygens were chosen; *d*_PEG_ is defined as the average of *Z*_MAX_ – *Z*_MIN_ of PEG oxygens. In addition, the PEG-chain behavior was analyzed by the fraction of PEG-chains involved in inter-chain interactions and the number of PEG units per chain interacting with phospholipids and CHOLs. Two PEG-chains are defined to be in contact if the minimum distance between heavy atom pairs in these chains is less than 4.5 Å. The contact between a PEG unit in a given chain and the membrane is defined by the same distance criterion.

## RESULTS AND DISCUSSION

### Illustration 1: Case 1 and Case 2

Our ionizable lipid models and system building are first illustrated by comparing our results with those from Ramezanpour et al^9^ (Case 1 in **Table 1**) and Ermilova and Swenson^10^ (Case 2). Consistent with Ramezanpour et al, neutral KC2 molecules accumulate in the bilayer center, whereas cationic head groups in KC2H stay in the bilayer-water interface (**Fig. 4** and **Figs. S2**a and **S3**a in ESI†). The extent of accumulation increases with KC2 concentration, which also increases the membrane thickness (**Fig. 5a**). In the presence of 45 mol% CHOL (KC2-PO-CH), the accumulation of KC2 is further promoted.

**Fig. 4.**
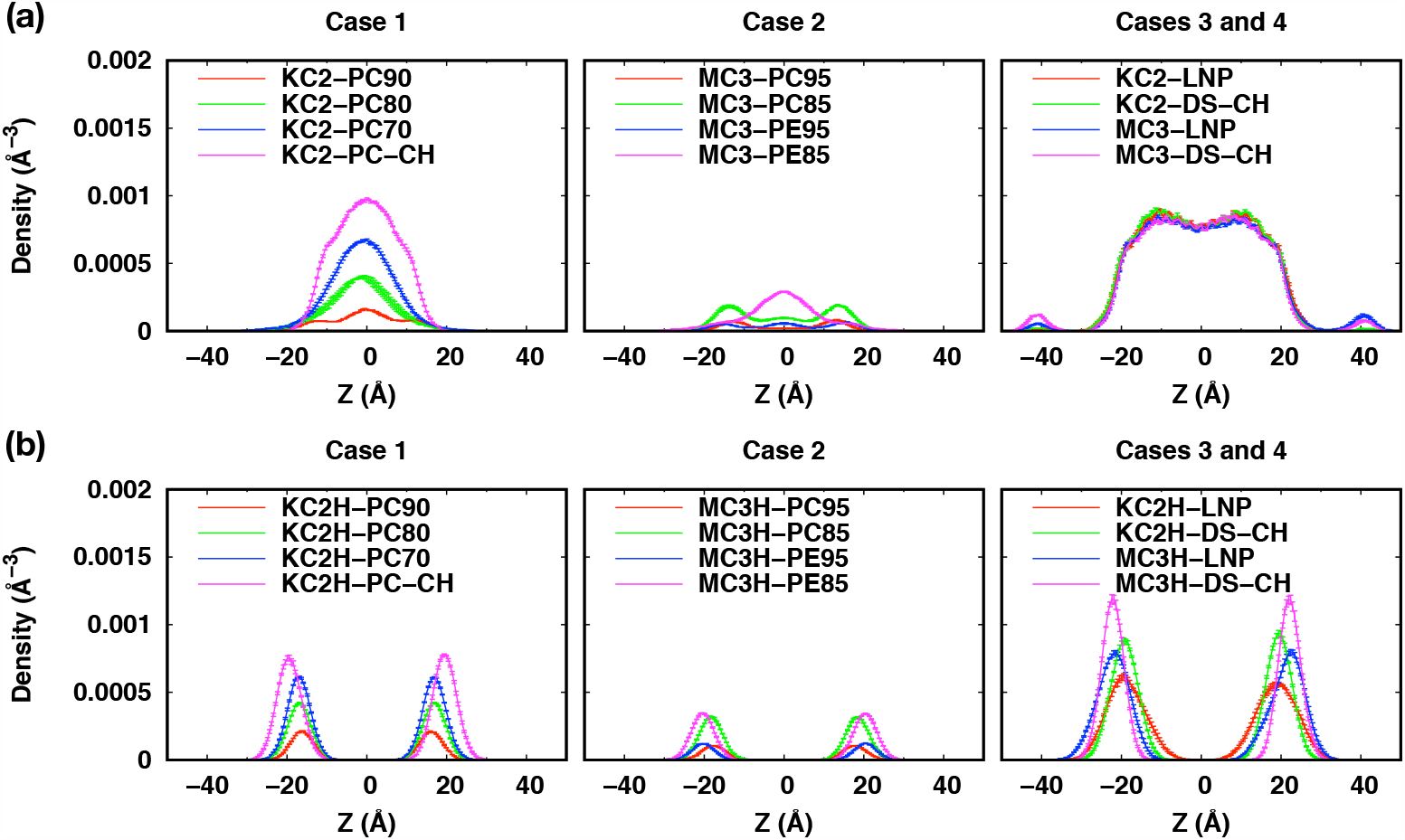
Density distributions of (a) neutral and (b) cationic DMA head groups. The profiles were calculated over three blocks from the last 600-ns trajectories. The error bars are the standard errors over the three blocks and they are too small to be seen clearly.

**Fig. 5.**
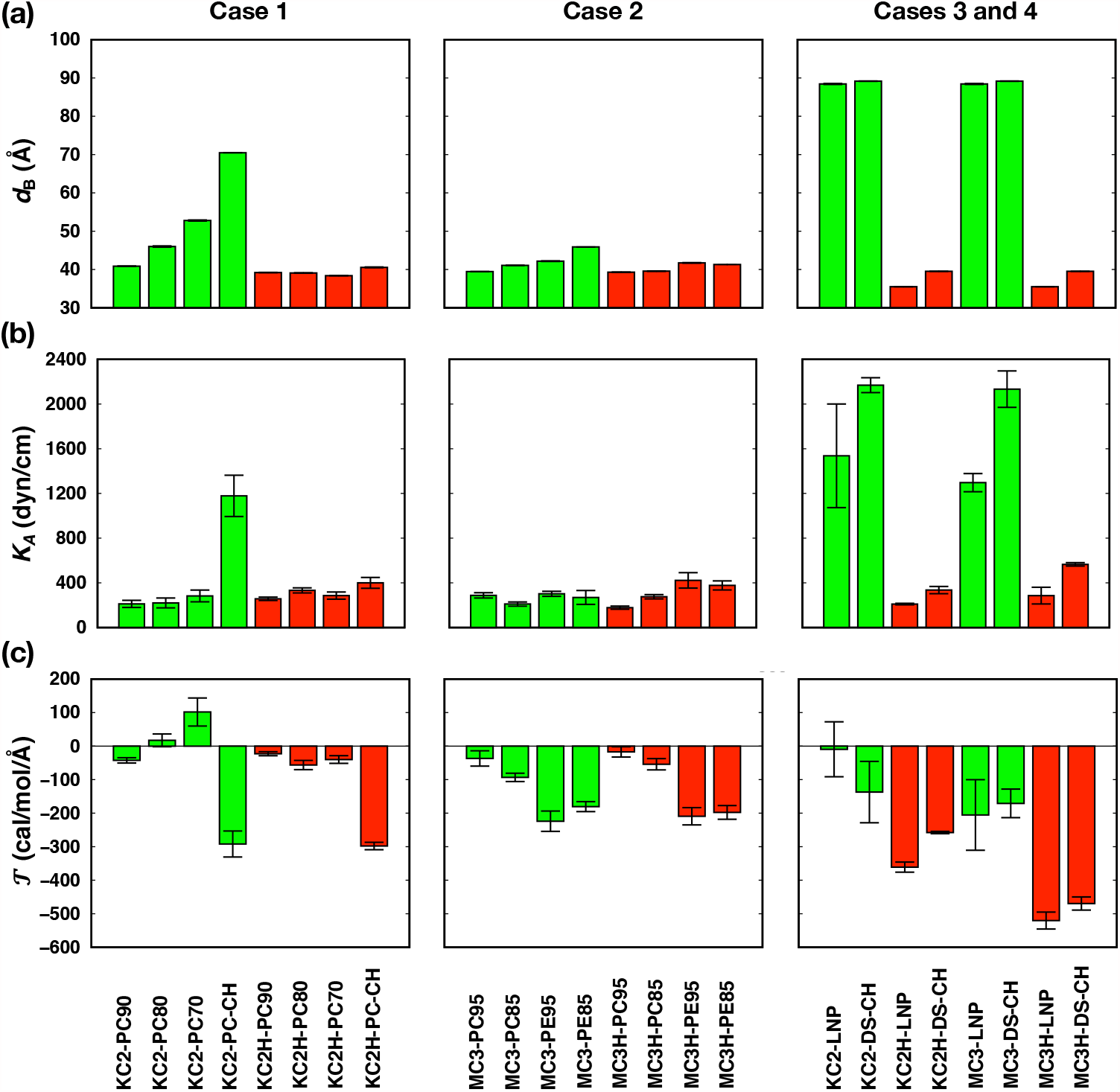
Bilayer properties: (a) membrane thickness (*d*_B_), (b) area compressibility modulus (*K*_A_), and (c) torque density (*𝒯*). The system names are given at the bottom of each column. Data for systems with neutral KC2 and MC3 lipids are shown in green bars and those for systems with cationic KC2H and MC3H lipids are shown in red bars. The averages were calculated over three 200-ns blocks from the last 600-ns data and the error bars represent the standard errors over the three blocks. These results are also summarized in **Table S1** in ESI†.

Observed POPC order parameters (*S*_CD_ in **Eq. 1**) in binary bilayers (without CHOL) are also consistent with Ramezanpour et al^9^ (at *T* = 313 K) in that *S*_CD_ are not sensitive to KC2 or KC2H concentrations (**Fig. S4**a in ESI†). *S*_CD_ of KC2’s all tail carbons remain low and are not sensitive to KC2 concentration, indicating that KC2 are randomly oriented in the bilayer center. KC2H’s *S*_CD_ are larger in the upper half of the tail, consistent with the fact that the cationic head groups stay in the bilayer-water interface. Due to the presence of two cis double bonds (**Fig. 1**), KC2H’s *S*_CD_ become smaller and comparable to those of KC2 in the lower half of its tail.

In the presence of 45% CHOL, POPC’s *S*_CD_ become significantly higher than the bilayers without CHOL, which can be attributed to the condensing effect of CHOL.^38–41^ The increase in POPC’s *S*_CD_ is less pronounced in KC2H-DS-CH (compared to those in KC2-DS-CH), indicating that repulsive interactions between cationic KC2H molecules reduce CHOL condensing effects. Expectedly, KC2’s *S*_CD_ are not changed (as KC2 molecules randomly accumulate in the bilayer center), while KC2H *S*_CD_ are significantly increased with 45% CHOL.

The above behavior of *S*_CD_ is well reflected in the calculated area compressibility modulus (*K*_A_) (**Fig. 5**b). *K*_A_ is not sensitive to KC2 concentration in bilayers without CHOL, and *K*_A_ shows slight increase at higher KC2 concentration. The opposing forces between cationic KC2H-KC2H repulsive interactions and CHOL condensing effects are also clearly shown in the resulting much less-increased *K*_A_ of KC2H-DS-CH compared to that of KC2-DS-CH. The observed *S*_CD_ and *K*_A_ are also consistent with the results from a recent simulation study of archaeal membranes,^42^ where concentration of menaquinone (accumulated in the bilayer center) does not significantly alter *K*_A_ and *S*_CD_ of the archaeal bilayers.

Although the extent of accumulation is less significant compared to KC2, neutral MC3 molecules also accumulate in the bilayer center, while MC3H head groups stay in the bilayer water-interface (**Fig. 4** and **Figs. S2**b and **S3**b in ESI†). The extents of MC3 accumulation in MC3/POPC binary bilayers are less pronounced than that of neutral KC2 in KC2/POPC binary bilayers, whereas the MC3 accumulation at 15 mol% MC3 in MC3/POPE bilayer (MC3-PE85) is comparable to KC2 accumulation in KC2/POPC bilayers (in between those in KC2-PC90 and KC2-PC80). The observed less pronounced accumulation of MC3 than KC2 can be attributed to MC3’s structure, where carbonyl oxygen atoms in MC3 head group can interact with the PE head group or glycerol backbone in PC/PE, which can hinder MC3 accumulation in the bilayer center (Ermilova and Swenson^10^ and **Fig. 1**)

The accumulation of MC3 was not observed in the previous study of Ermilova and Swenson^10^ with MC3/DOPC and MC3/DOPE binary bilayers. We attribute the observed MC3 accumulation in Case 2 to the difference in the force fields between our work and theirs. The more pronounced accumulation of MC3 in MC3/POPE bilayers than in MC3/POPC bilayers can also be attributed to a negative spontaneous curvature of POPE and a smaller spontaneous curvature of POPC,^43^ which could enable POPE monolayers to better accommodate domains of neutral MC3 in the bilayer center.

POPE’s S_CD_ are slightly higher than POPC’s *S*_CD_ (**Fig. S4**b in ESI†), which is consistent with the findings from Ermilova and Swenson^10^ in that the carbonyl oxygen atoms in MC3 can form additional hydrogen bonds with PE amine hydrogen atoms. Similar to Case 1, phospholipids’ *S*_CD_ are not sensitive to MC3 or MC3H concentration. *S*_CD_ of MC3 and MC3H behave similarly to phospholipids’ *S*_CD_ except low MC3’s *S*_CD_ in MC3-PE85 (due to MC3 aggregation in the bilayer center). Consistent with *S*_CD_, *K*_A_ are not sensitive to MC3 and MC3H concentration, and *K*_A_ from binary bilayers with POPE are slightly larger than those from binary bilayers with POPC.

### Illustration 2: Case 3 and Case 4

As a second illustration, we now focus on more realistic LNP membranes with or without PEG-lipids. Consistent with Case 1 and Case 2, KC2 and MC3 accumulate in the bilayer center, and KC2H and MC3H head groups stay in the bilayer-water interface (**Figs. 4, 6**, and **S5** (ESI†)). Due to accumulated KC2 and MC3, the membrane thickness increases significantly compared to the mixed bilayers with KC2H or MC3H (**Fig. 5a**).

**Fig. 6.**
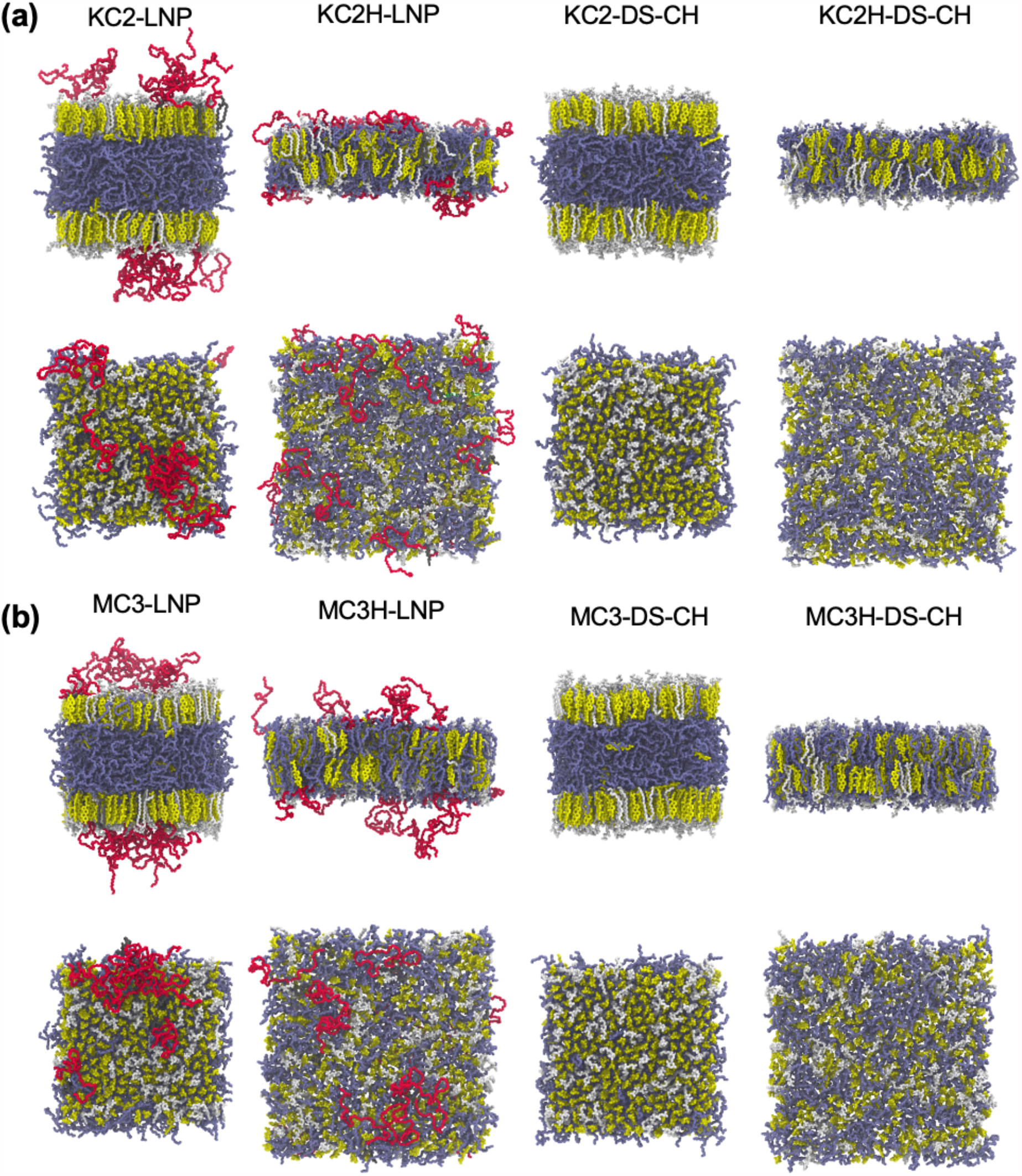
Final snapshots of (a) Case 3 and (b) Case 4 systems. Each system name is given on top of each column showing the side and top views. The color code is the same as in **Fig. 3**.

As expected, the surface areas are comparable between KC2-and MC3-containing bilayers as KC2s and MC3s are segregated to the bilayer center (**Fig. S1**c in ESI†). The surface areas of KC2H-containg bilayers are consistently larger than those of MC3H-containing bilayers. This can be attributed to the difference in head group structures between KC2H and MC3H (**Fig. 1**), where a bulkier five membered ring in KC2H compared to the carbonyl group in MC3H makes lipid packing less tight. This also results in a higher Cl^-^ density due to higher surface charge density in MC3H-containing bilayers (**Figs. 4, 7**, and **S5** (ESI†)).

**Fig. 7.**
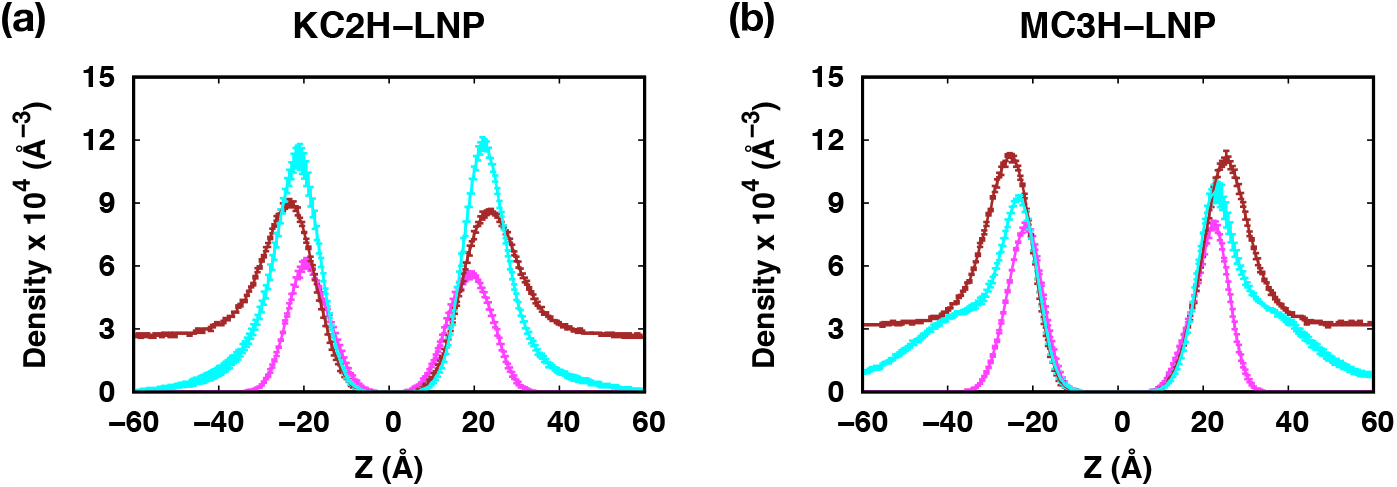
Density profiles of ionizable lipids, PEG-chains, and Cl^-^ions along the membrane normal (i.e., the Z axis) for (a) KC2H-LNP and (b) MC3H-LNP. Cl^-^densities are scaled up by a factor of 2 for easier comparison between panels. The profiles were calculated over three 200-ns blocks from the last 600-ns trajectories. The error bars are the standard errors over the three blocks and they are too small to be seen clearly. Each component is shown in different colors: nitrogen in ionizable lipids (magenta), oxygen in PEG-chains (cyan), and Cl^-^ions (brown).

Lateral lipid packing in Case 3 and Case 4 is consistent with Case 1 and Case 2. *S*_CD_ of DSPC and PEG-lipid are larger for mixed bilayers with neutral KC2/MC3 than those with KC2H/MC3H (**Fig. S7** in ESI†) due to the counteraction between cationic KC2H/MC3H repulsive interactions and CHOL’s condensing effect. *S*_CD_ of KC2/MC3 remain low due to their accumulation at the bilayer center and those of KC2H/MC3H are significantly higher. The calculated *K*_A_ are consistent with the observed *S*_CD_, where *K*_A_ of KC2H/MC3H-containing membranes significantly smaller than those of KC2/MC3-containing membranes (**Fig. 5b**).

The presence of PEG-lipids slightly reduces *S*_CD_ of DSPC and KC2H, which can be attributed to shorter myristoyl (14:0) tails in PEG-lipids that induce less order than stearoyl (18:0) tails in DSPC. The decrease in *K*_A_ with PEG-lipids is consistent with the observed behavior of DSPC’s *S*_CD_. With PEG-lipids, *K*_A_ of LNPs with cationic lipids (KC2H or MC3H) fit into typical *K*_A_ of fluid-phase membranes.^43,44^

### Effects of ionizable cationic lipids on PEG-membrane interactions

PEG-chains in our simulations are flexible (**Movies S1**-**4** in ESI†). The PEG thickness (*d*_PEG_) is comparable or smaller than the end-to-end distance, ⟨*h*^2^⟩^1/2^ (**Fig. S6** in ESI†), indicating that PEG-chains are in the mushroom conformation regime (expected from low surface density).^8,45,46^ PEG-chains in KC2-LNP and MC3-LNP frequently make inter-PEG contacts, while those in KC2H-LNP and MC3H-LNP interact more with the membrane (**Fig. S7** in ESI†).

More frequent inter-PEG contacts in LNPs with KC2 or MC3 than those in LNPs with KC2H or MC3H are consistent with the increased surface PEG-lipid concentration from 2% to ∼3.3% due to the accumulation of KC2 and MC3 in the bilayer center. The attractive interactions between cationic KC2H head groups and PEG oxygens are responsible for the observed PEG-membrane interactions, which also makes the PEG conformations extended more preferentially on the membrane surface (**Fig. 6 and S6**).

The interactions between MC3H head groups and PEG oxygen atoms become weaker in MC3H-LNP, due to a higher concentration of Cl^-^ ions near the membrane (**Fig. 7**). As a result, the PEG-membrane contacts becomes smaller than those in KC2H-LNP (**Fig. S7**b) and the PEG end-to-end distance becomes isotropic (**Fig. S6**b).

### Effects of ionizable cationic lipids and PEG-lipids on membrane curvature

Inclusion of ionizable cationic lipids and/or PEG-lipids affects lateral packing of membranes (see above), as well as the membrane curvature. These effects are reflected in the monolayer torque density, 𝒯 (see **Lateral pressure profile and torque density** in ESI†). The sign of 𝒯 dictates the direction of applied (bending) torque, and the sign of its change, *Δ*𝒯, between different concentrations of ionizable cationic lipids and/or PEG-lipids indicates the direction of the induced curvature. In this work, we adopt a typical sign convention of the curvature: a monolayer at a positive curvature is convex to the head group side, and vice versa. In this convention, positive *Δ*𝒯 between two systems indicates that a positive curvature is induced, and vice versa.

At the lowest KC2/KC2H concentration (10 mol%), 𝒯 are small negative numbers (**Fig. 5c**) close to that of pure POPC bilayer (around -30 cal/mol/Å)^43^. 𝒯 increases with KC2 concentration, indicating that accumulated KC2 at the bilayer center induces positive curvature to the monolayers. However, 𝒯 are much less sensitive to KC2H concentration, indicating that KC2H do not strongly affect the membrane curvature. This can be attributed to the structurally similar head groups of KC2H to PC of phospholipids, which have small intrinsic curvature.^43^ For CHOL-containing bilayers, 𝒯 becomes noticeably negative compared to those of CHOL-free bilayers, consistent with the negative curvature induced by CHOL.^47^

Opposite to the 𝒯 variation in KC2/POPC bilayers, 𝒯 of MC3/POPC binary bilayers becomes more negative at a higher MC3 concentration (Case 2 in **Fig. 5c**), indicating that MC3 induces negative curvature with POPC, which is likely due to the interactions between carbonyl oxygen in MC3 and the glycerol backbone in POPC.^10^ Opposite to the 𝒯 variation in MC3/POPC, increasing MC3 concentration makes 𝒯 become more positive in MC3/POPE binary bilayer due to the accumulation of MC3 in the bilayer center, rendering the POPE monolayer curvature less negative and thus the entire system becomes more stable. Similarly to KC2H, 𝒯 are not sensitive to MC3H concentration both in MC3H/POPC and MD3H/POPE binary bilayers. Again, this can be attributed to similar head group structures between MC3H and PC.

Effects of PEG-lipids on 𝒯 of KC2-containing bilayers appear to be minimal, as shown by statistically similar 𝒯 between KC2-LNP and KC2-DS-CH. 𝒯 between MC3-LNP and MC3-DS-CH are also similar. These results indicate that the interactions between PEG-lipid and KC2/MC3 are likely to be weak (as KC2/MC3 are located in the bilayer center), and do not appear to affect the membrane curvature.

For KC2H-containing bilayers, the interactions between KC2H and PEG-lipid significantly affect 𝒯, as shown by noticeably more negative 𝒯 of KC2H-LNP compared to that of KC2H-DS-CH. Such induced negative curvature arises from the attractive interactions between KC2H and PEG oxygen. The same trend is observed between MC3H-LNP and MC3-DS-CH, though the extent of induced negative curvature is smaller, as a higher Cl^-^ ion concentration near the membrane due to higher surface charge density (**Fig. 7**) weakens the attractive interactions between MC3H and PEG oxygen.

### Implications to genetic drug delivery

It has been accepted that the incorporation of helper lipids including PEG-lipids in LNPs is required to increase the circulation time, so that LNPs can reach target cells.^5^ After their uptake to cytoplasm, the cationic lipids and endogenous anionic lipids produce a non-bilayer structure at low pH in the endosome, i.e., below the pKa of ionizable cationic lipids, which results in the disruption of the endosomal membrane and release of genetic drugs into the cytoplasm.^1^

Prior to the delivery process, PEG-chains need to be detached from LNPs for efficient endocytosis of nanocarriers loaded with genetic drugs. Otherwise, drug activity can be dramatically decreased by reduction of the cellular uptake of LNPs^48–50^ and accelerated blood clearance after repeated doses of PEGylated liposomes.^51,52^ The cleavable PEGylation^53–55^ is one of the approaches that can overcome this issue, i.e., the so-called PEG dilemma.

Our results show that PEG-lipid linkage can be exposed more in KC2H-containing LNPs than MC3H-containing ones (**Figs. 6** and **7**, and **Fig. S6**b and **Movies S1**-**4** in ESI†). For efficient cleavage of PEGylation, the linkage between PEG and lipid tails are desired to be exposed to extracellular or intracellular microenvironment, such as temperature, pH, specific enzyme, reductive conditions, etc.^56–58^ Therefore, our results implicate that one can tune cationic ionizable lipid-PEG interactions to achieve desired cleavage of PEGylation for rational LNP design and optimal drug delivery.

## CONCLUSIONS

We have described an important extension of CHARMM-GUI *Membrane Builder* to model and simulate LNPs with various (ionizable) cationic lipids and PEG-lipids. To illustrate this new feature, we have built and simulated all systems of two previous simulation studies, Ramezanpour et al.^9^ and Ermilova and Swenson.^10^ As another more practical illustration to demonstrate *Membrane Builder*’s capability to build a realistic LNP patch, we built and simulated KC2-and MC3-based LNP membranes with very high cholesterol and ionizable cationic lipids together with 2 mol% PEG-lipids.

In our simulations, cationic lipids (KC2H and MC3H) stay at the bilayer-water interface, and neutral KC2 and MC3 molecules segregate into the bilayer center, though MC3 accumulation is less pronounced. The accumulation of KC2/MC3 does not alter lateral lipid packing, which has been also reported in a recent study of archaeal membranes.^42^ The results from binary bilayers of KC2/KC2H and POPC are consistent with Ramezanpour et al. The less pronounced MC3 accumulation is consistent with observed MC3 and phospholipid interactions in Ermilova and Swenson. However, MC3 accumulation was not observed in Ermilova and Swenson (binary bilayers of MC3 and DOPC/DOPE), which is attributed to differences in the force fields between our work and theirs.

PEG-chains are flexible and frequently extended to bulk water. They are extended more preferentially on the membrane surface of LNP membranes with cationic lipids due to the interactions between PEG oxygen and their cationic head groups. The presence of PEG-lipids appears to relax lateral lipid packing in LNP membranes by decreasing *K*_A_ and *S*_CD_ of DSPC and cationic lipids (KC2H and MC3H). Interestingly, *K*_A_ of LNP membranes with cationic lipids (KC2H or MC3H) fit into typical *K*_A_ of fluid-phase membranes. Our results show that the interactions between PEG oxygen and cationic lipid head groups induce a negative curvature. Our results also show that PEG-cationic ionizable lipid interactions can be tuned for the desired cleavage of PEGylation for optimal drug delivery. Therefore, it would be an interesting future study to model and simulate LNPs with encapsulated nucleic acids in different environments.

## ACKNOWLEDGEMENTS

This study was supported in part by grants from NIH GM138472 and NSF MCB-1810695.

†Electronic supplementary information (ESI) available: Additional section for lateral pressure profile and torque density; table showing membrane thickness, area compressibility modulus, and torque density; figures showing time series of area, snapshots, density profiles, *S*_CD_ order parameters, end to end distance and thickness of PEG, fraction of PEG-lipid interacting with other PEG-lipids, and the number of PEG units per chain interacting with membrane; movies for Cases 3 and 4.

## Notes

### Competing Interest Statement

The authors have declared no competing interest.

