## Supporting Information for "CHARMM-GUI Membrane Builder for Lipid Nanoparticles with Ionizable Cationic Lipids and PEGylated Lipids"

**Lateral pressure profile and torque density**

The lateral pressure profile, $p(Z) = p_T(Z) - p_N(Z)$, of a membrane system describes non-uniform interactions between its components along the membrane normal (i.e., the Z-axis) that arises from their heterogeneous distributions. Here, $p_T(Z) = [p_{xx}(Z)+p_{yy}(Z)]/2$ and $p_N(Z) = p_{zz}(Z) = p_N$ are the tangential and normal components of pressure tensors to the membrane surface. The first moment of $p(Z)$ is the (bending) torque density, $\mathcal{T}$, applied to the membrane. For the upper leaflet, it is given by

$$\mathcal{T} = \int_0^{L_z/2} dZ\, Z\, p(Z) \quad \text{(S1a)}$$

where $L_z$ is the system size along the membrane normal and $Z = 0$ is the bilayer center. For a stress-free symmetric bilayer, the monolayer torque density can be connected to its bending modulus ($k_c$) and its spontaneous curvature ($c_0$) as $\mathcal{T} = k_c c_0$. In a typical monolayer curvature convention, a monolayer at a positive curvature is convex to the head group side, and vice versa (see **Figure S0**). Positive or negative $\mathcal{T}$ for a monolayer implies that there exists a torque, which would increase or decrease its curvature, respectively. A consistent $\mathcal{T}$ for the lower leaflet with this sign convention is given by

$$\mathcal{T} = \int_0^{-L_z/2} dZ\, Z\, p(Z) \quad \text{(S1b)}$$

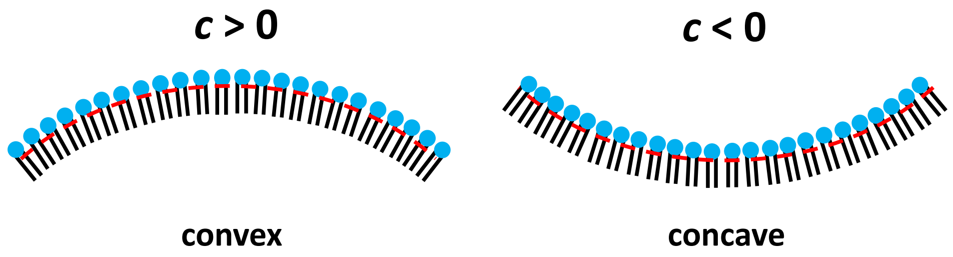


**Figure S0.** Typical monolayer curvature ($c$) convention. Head groups are shown as blue circles and tails are shown as black lines.

**Table S1**. Membrane thickness ($d_B$), area compressibility modulus ($K_A$), and torque density ($\mathcal{T}$).[a]

| Case 1 | $d_B$ (Å) | $K_A$ (dyn/cm) | $\mathcal{T}$ (cal/mol/Å) |
|---|---|---|---|
| KC2-PC90 | 40.9 (0.0) | 211.6 (31.7) | –43 (8) |
| KC2-PC80 | 46.0 (0.2) | 220.3 (44.1) | 17 (19) |
| KC2-PC70 | 52.8 (0.1) | 282.8 (52.6) | 102 (42) |
| KC2-PC-CH | 70.5 (0.0) | 1178.3 (184.9) | –292 (39) |
| KC2H-PC90 | 39.2 (0.0) | 275.3 (14.9) | –23 (6) |
| KC2H-PC80 | 39.1 (0.0) | 332.6 (22.8) | –56 (14) |
| KC2H-PC70 | 38.4 (0.0) | 285.8 (32.3) | –40 (11) |
| KC2H-PC-CH | 40.6 (0.1) | 400.0 (48.5) | –298 (11) |
| **Case 2** | $d_B$ (Å) | $K_A$ (dyn/cm) | $\mathcal{T}$ (cal/mol/Å) |
| MC3-PC95 | 39.5 (0.0) | 287.9 (23.8) | –37 (23) |
| MC3-PC85 | 41.1 (0.0) | 210.4 (18.9) | –93 (12) |
| MC3-PE95 | 42.1 (0.1) | 302.2 (22.5) | –224 (30) |
| MC3-PE85 | 45.9 (0.0) | 269.3 (62.3) | –181 (15) |
| MC3H-PC95 | 39.3 (0.0) | 177.8 (14.2) | –17 (15) |
| MC3H-PC85 | 39.6 (0.0) | 275.8 (19.1) | –54 (17) |
| MC3H-PE95 | 41.7 (0.1) | 422.1 (69.0) | –209 (26) |
| MC3H-PE85 | 41.3 (0.0) | 377.1 (40.6) | –198 (20) |
| **Case 3** | $d_B$ (Å) | $K_A$ (dyn/cm) | $\mathcal{T}$ (cal/mol/Å) |
| KC2-LNP | 88.3 (0.1) | 1880.8 (507.8) | –10 (82) |
| KC2-DS-CH | 89.2 (0.0) | 2168.9 (66.7) | –137 (92) |
| KC2H-LNP | 35.5 (0.0) | 210.0 (6.9) | –361 (15) |
| KC2H-DS-CH | 39.5 (0.0) | 355.5 (32.2) | –258 (3) |
| **Case 4** | $d_B$ (Å) | $K_A$ (dyn/cm) | $\mathcal{T}$ (cal/mol/Å) |
| MC3-LNP | 87.7 (0.1) | 1296.6 (81.4) | –205 (105) |
| MC3-DS-CH | 87.9 (0.2) | 2133.3 (163.0) | –171 (43) |
| MC3H-LNP | 41.4 (0.1) | 285.7 (74.8) | –520 (25) |
| MC3H-DS-CH | 43.8 (0.1) | 565.3 (15.5) | –470 (20) |

[a]The averages were calculated over three blocks from the last 600-ns trajectories. The standard errors over three 200-ns blocks are given in parentheses.

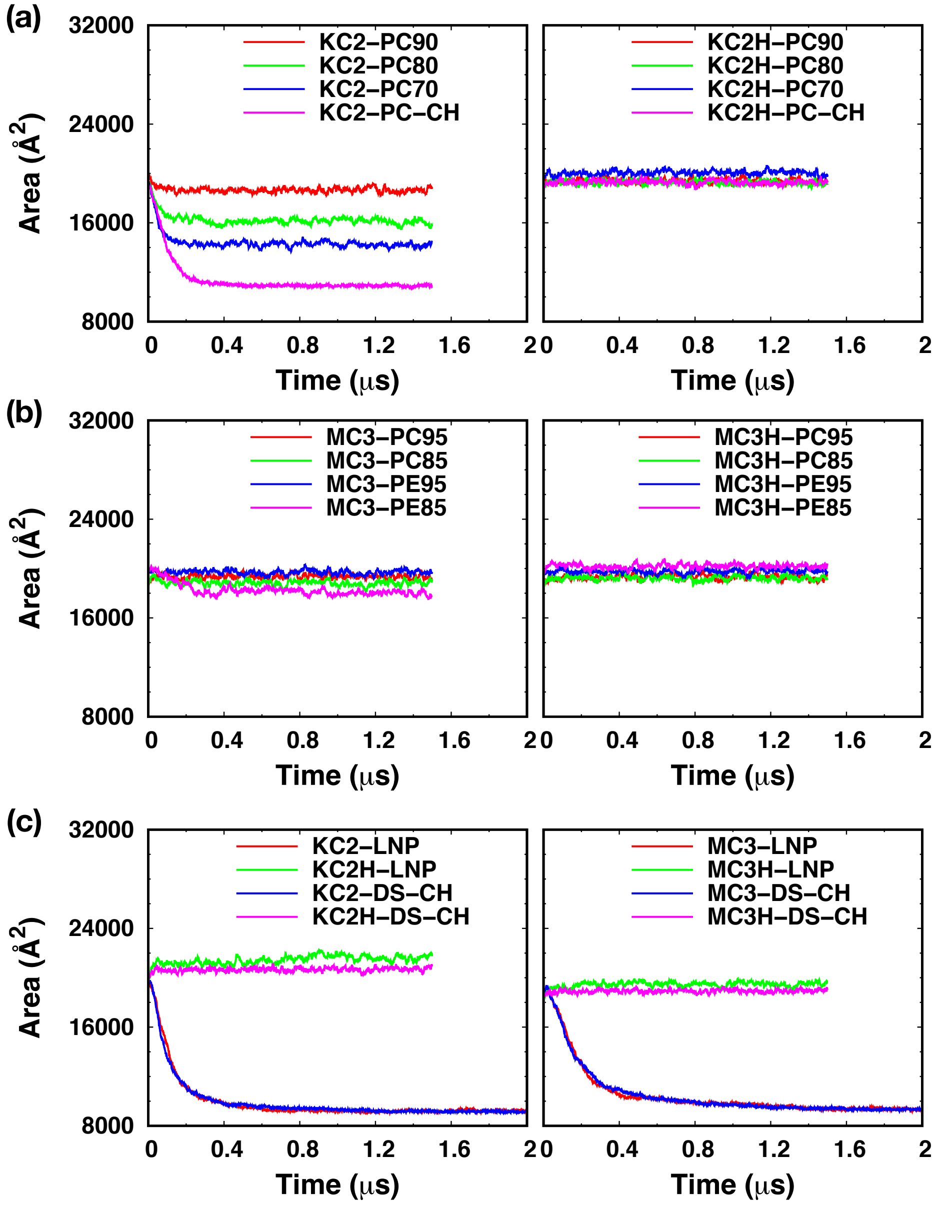


**Figure S1.** Time series of the XY membrane area of (a) Case 1, (b) Case 2, and (c) Cases 3 and 4 systems in **Table 1**.

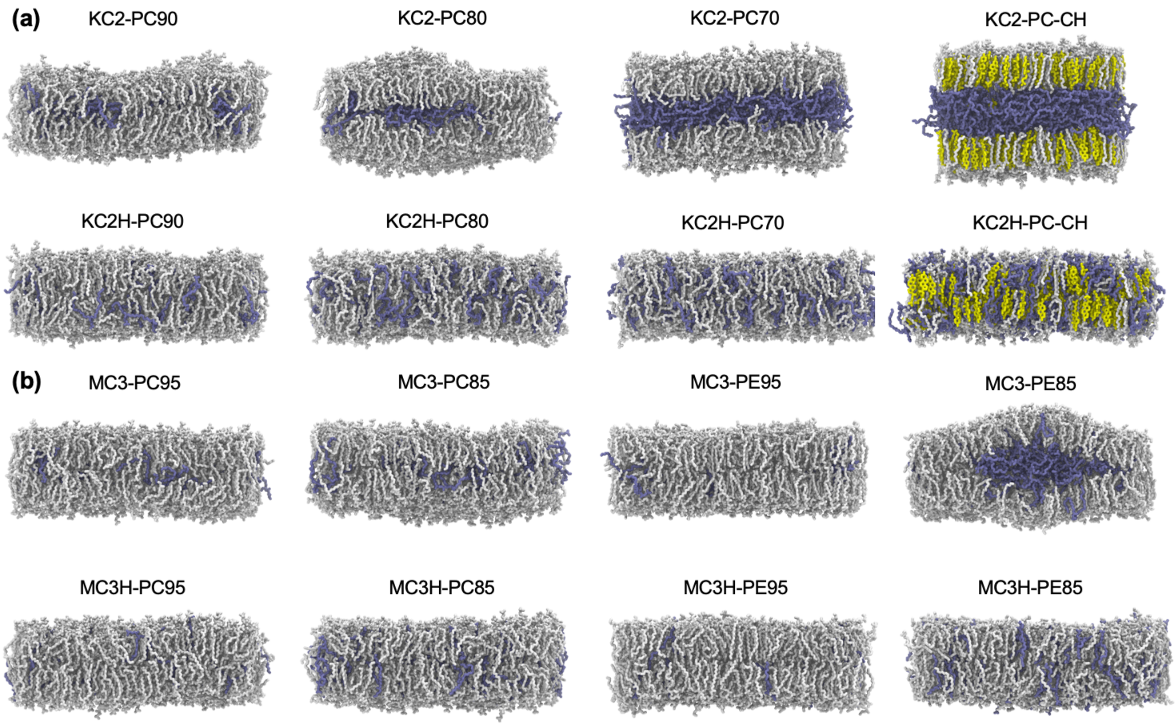


**Figure S2**. Final snapshots of (a) Case 1 and (b) Case 2 systems. Each system name is shown on top of each panel. The color code is the same as in **Figure 3**.

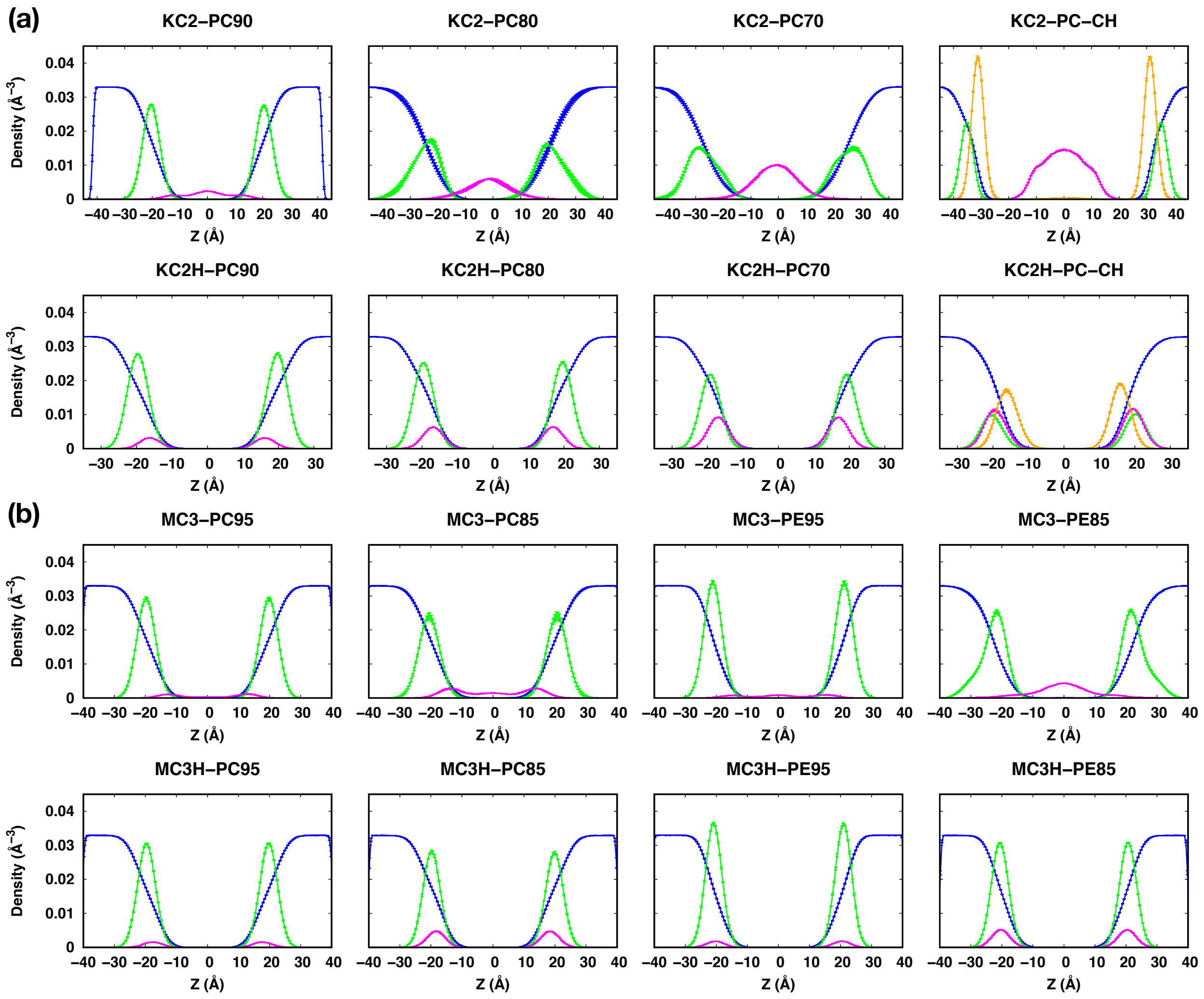


**Figure S3**. Component density profiles along the membrane normal (i.e., the *Z* axis) in (a) Case 1 and (b) Case 2 systems. Each system name is shown on top of each panel. The profiles were calculated over three blocks from the last 600-ns trajectories. The error bars are the standard errors over the three blocks and they are too small to be seen clearly. Each component is shown in different colors: water oxygen (blue), phosphorous in lipid head group (green), nitrogen in ionizable lipids (magenta), and CHOL oxygen (orange). For better visibility, densities are scaled up by a factor of 15 except water.

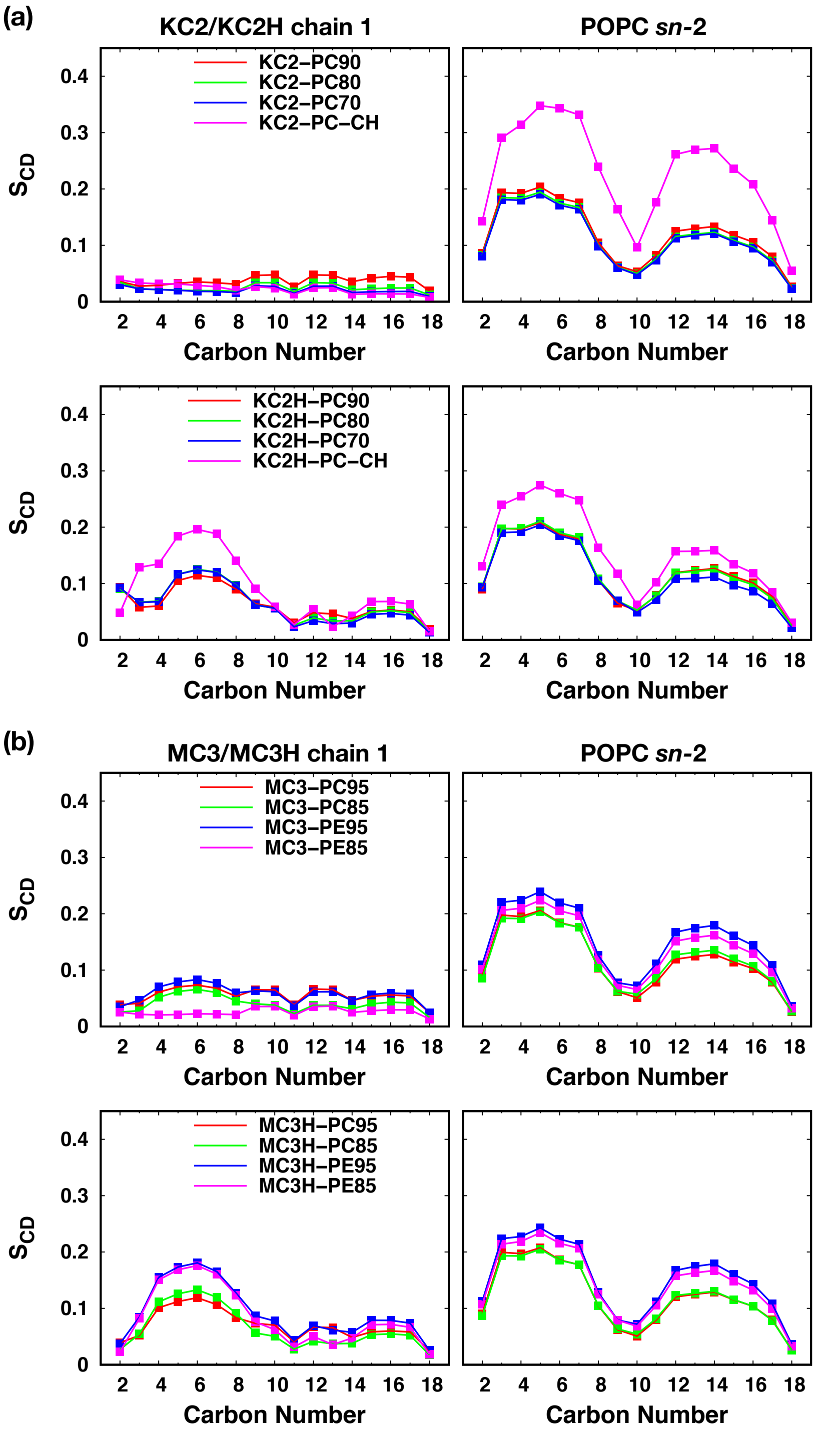


**Figure S4.** $S_{CD}$ order parameters of (a) Case 1 and (b) Case 2 systems. $S_{CD}$ of ionizable lipids are shown in the left column and those of phospholipids are shown in right column. The lipid and chain names are shown on top of each column and the system names are shown in the left panels. The average $S_{CD}$ were calculated over three blocks from the last 600-ns trajectories. The error bars are the standard errors over the three blocks and they are too small to be seen clearly.

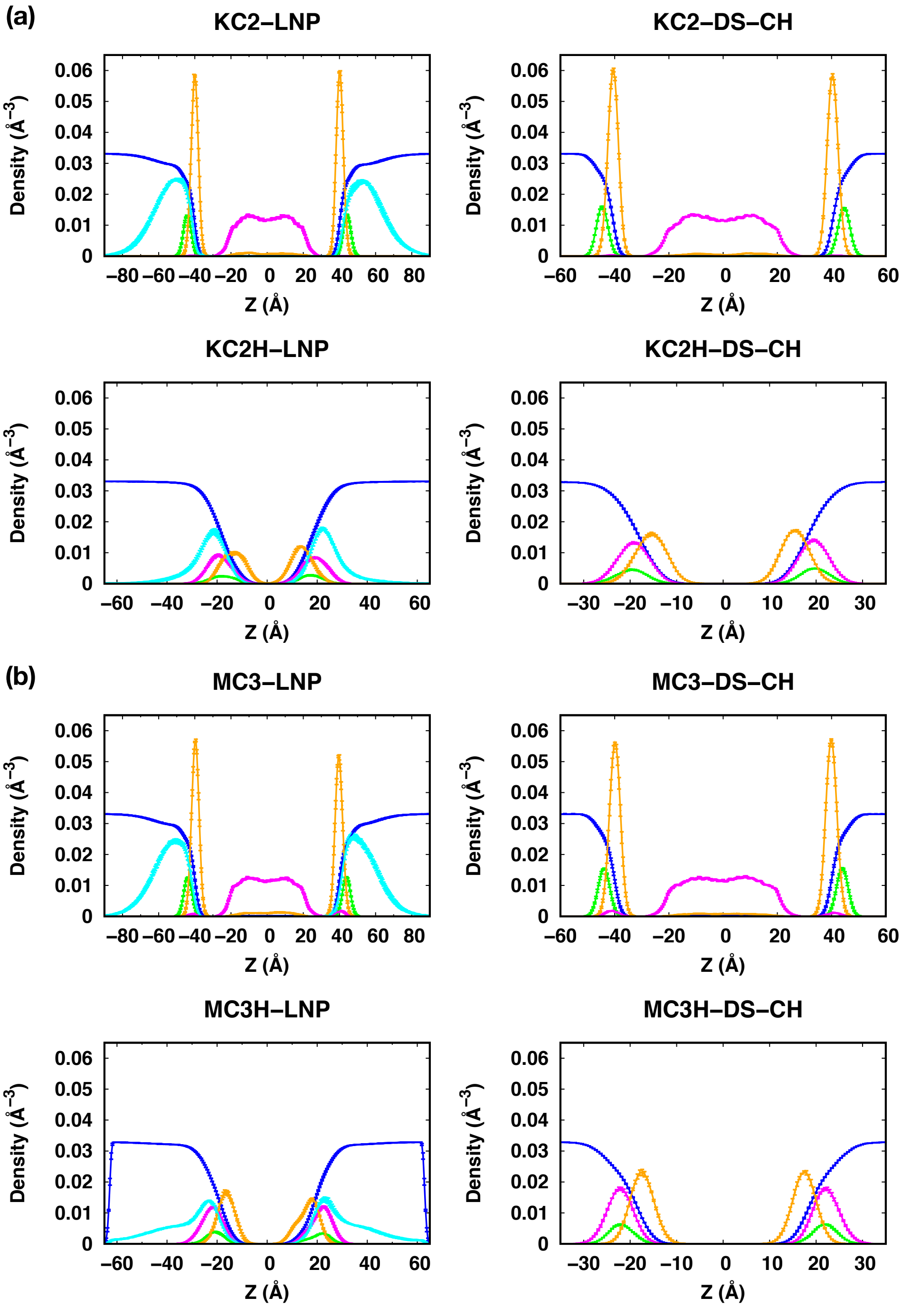


**Figure S5**. Component density profiles along the membrane normal (i.e., the Z axis) in (a) Case 3 and (b) Case 4 systems. Each system name is shown on top of each panel. The profiles were calculated over three blocks from the last 600-ns trajectories. The error bars are the standard errors over the three blocks and they are too small to be seen clearly. Each component is shown in different colors: water oxygen (blue), phosphorous in lipid head group (green), nitrogen in ionizable lipids (magenta), and CHOL oxygen (orange), and PEG chain (cyan). For better visibility, densities are scaled up by a factor of 15 except water.

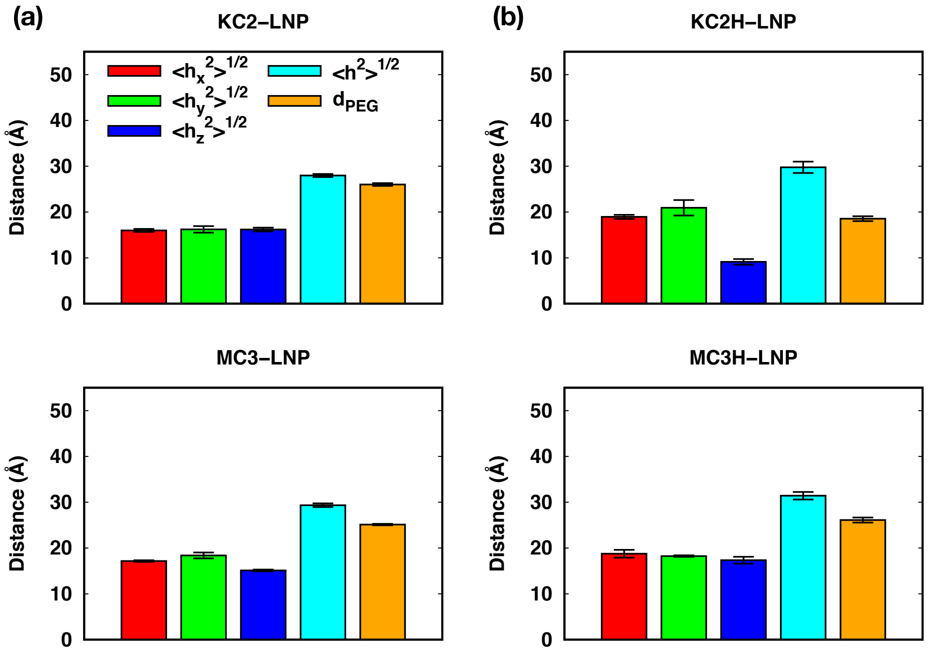


**Figure S6.** The end-to-end distance, $\langle h^2\rangle^{1/2}$, and the PEG thickness, $d_{PEG}$, for (a) LNP with neutral ionizable lipids (left) and (b) LNP with cationic ionizable lipids (right). $\langle h^2\rangle^{1/2}$ and $d_{PEG}$ are shown in cyan and orange bars, respectively. Shown together are *X*-, *Y*-, and *Z*-components of $\langle h^2\rangle^{1/2}$, $\langle h_x^2\rangle^{1/2}$, and $\langle h_y^2\rangle^{1/2}$, $\langle h_z^2\rangle^{1/2}$ (red, green, and blue bars, respectively). The system names are given on top of each panel. The averages were calculated over three 200-ns blocks from the last 600-ns data and the error bars represent the standard errors over the three blocks.

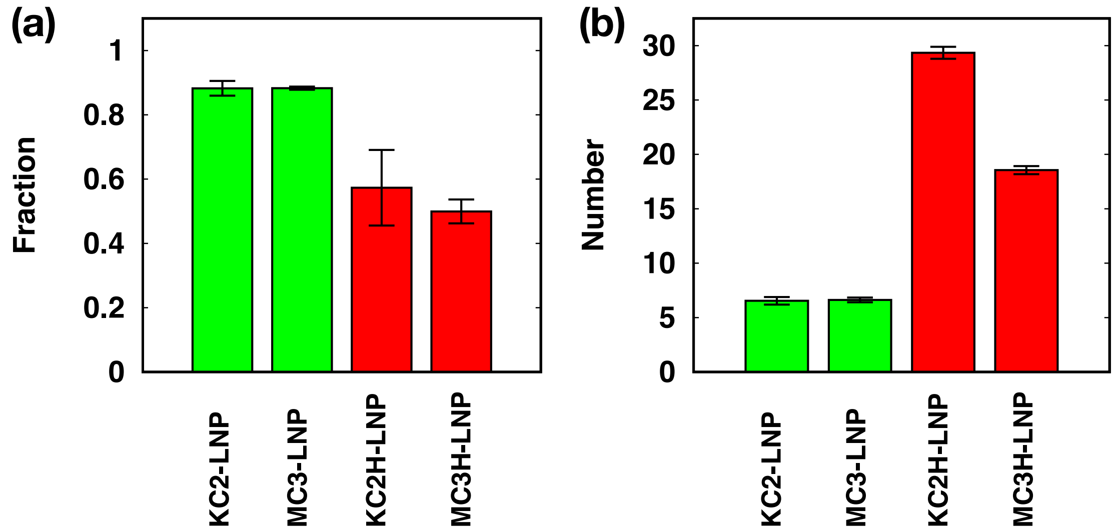


**Figure. S7** PEG-PEG and PEG-membrane interactions in LNP bilayers: (a) fraction of PEG-lipids having inter-PEG interactions and (b) number of PEG units per PEG-chain interacting with membrane. The system names are given at the bottom of each panel. Data for systems with neutral ionizable lipids are shown in green bars and those with cationic ionizable lipids are shown in red bars. The averages were calculated over three 200-ns blocks from the last 600-ns data and the error bars represent the standard errors over the three blocks.

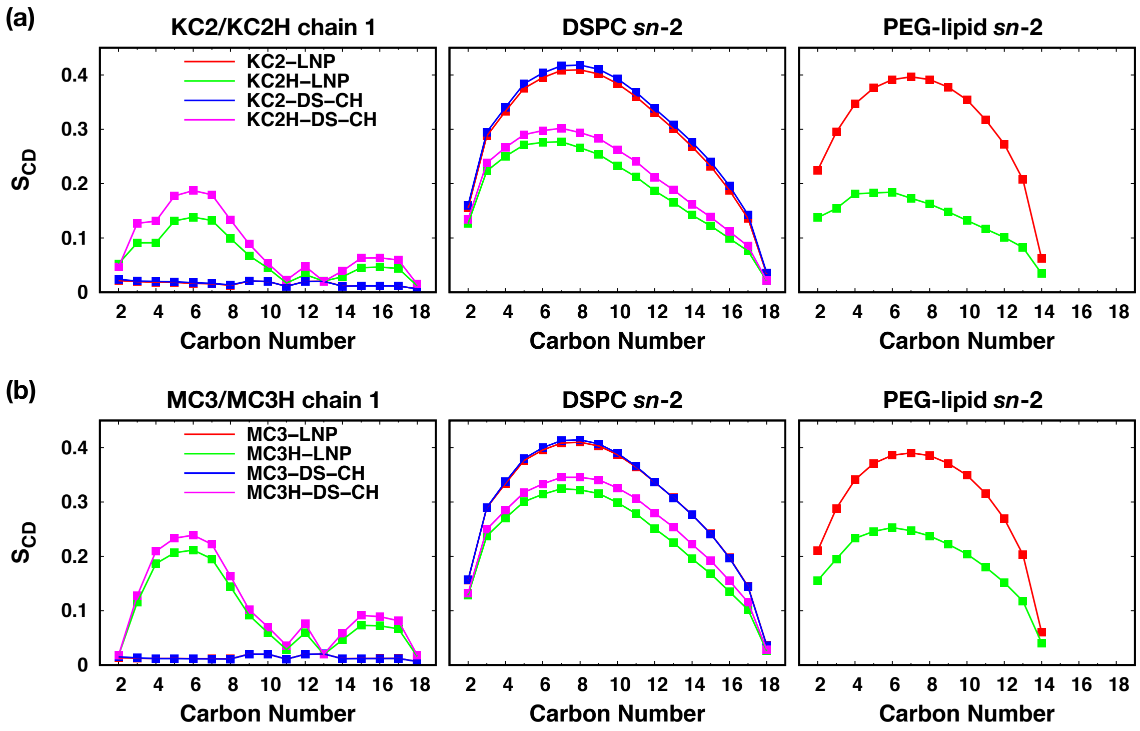


**Figure S8.** $S_{CD}$ order parameters of (a) Case 3 and (b) Case 4 systems. $S_{CD}$ of ionizable lipids, DSPC, and PEG-lipids are shown in the left, middle, and right columns, respectively. The lipid and chain names are shown on top of each column and the system names are shown in the left panels. The average $S_{CD}$ were calculated over three blocks from the last 600-ns trajectories. The error bars are the standard errors over the three blocks and they are too small to be seen clearly.

**Movie S1**. Movie for KC2-LNP with top and side views. The color code is the same as in **Fig.3**.

**Movie S2**. Movie for KC2H-LNP with top and side views. The color code is the same as in **Fig.3**.

**Movie S3**. Movie for MC3-LNP with top and side views. The color code is the same as in **Fig.3**.

**Movie S4**. Movie for MC3H-LNP with top and side views. The color code is the same as in **Fig.3**.
